# Influence of *APOA5* locus on the treatment efficacy of three statins: evidence from a randomized pilot study in Chinese subjects

**DOI:** 10.1101/213629

**Authors:** Sha Hua, Chuanxiang Ma, Jun Zhang, Jing Li, Weiwei Wu, Ning Xu, Guanghua Luo, Jianrong Zhao

**Affiliations:** Department of Cardiology, Ruijin Hospital Luwan Branch, School of Medicine, Shanghai Jiao Tong University, South Chongqing Rd. No.149, Shanghai 200020, China; Department of Pathology, The Affiliated Hospital of Weifang Medical University, Weifang 261000, China; Comprehensive Laboratory, Third Affiliated Hospital of Soochow University, 185 Juqian Street, Changzhou 213003, China; Section of Clinical Chemistry and Pharmacology, Institute of Laboratory Medicine, Lund University, S-221 85 Lund, Sweden

**Keywords:** *APOA5* genotype, statins, triglycerides, dyslipidemia

## Abstract

Pharmacogenetics or pharmacogenomics approaches are important for addressing the individual variabilities of drug efficacy especially in the era of precision medicine. One particular interesting gene to investigate is *APOA5* which has been repeatedly linked with the inter-individual variations of serum triglycerides. Here, we explored *APOA5*-statin interactions in 195 Chinese subjects randomized to rosuvastatin (5-10 mg/day), atorvastatin (10-20 mg/day), or simvastatin (40 mg/day) for 12 weeks by performing a targeted genotyping analysis of the *APOA5* promoter SNP rs662799 (-1131T>C). There were no significant differences between the treatment arms for any of the statin-induced changes in clinical biomarkers. Reductions in LDL cholesterol were influenced by the *APOA5* genotype in all three treatment groups. By contrast, changes in HDL cholesterol and triglycerides were only affected by the *APOA5* genotype in the atorvastatin and simvastatin groups and not in the rosuvastatin group. Our results support earlier findings indicating that rosuvastatin is a better treatment option and that future studies should consider stratifying subjects not only by genetic background but also by statin type.

**Abbreviations:**

ApoA5: apolipoprotein A5
BMI: body mass index
FFA: free fatty acids
HDLc: high-density lipoprotein cholesterol
LDLc: low-density lipoprotein cholesterol
Lp(a): lipoprotein(a)
SNP: single nucleotide polymorphism
T2D: type 2 diabetes
Tc: total cholesterol
Tg: triglycerides.

## Introduction

Although statins are the most prescribed class of drugs worldwide for prevention of various cardiovascular diseases, about one third of patients do not respond well to this therapy with respect to the lipid-lowering effect, suggesting that pharmacogenomics (Postmus et al., 2014) or other environmental factors such as diet (Jenkins et al., 2005) or the gut microbiome (Kaddurah-Daouk et al., 2011) may play substantial roles. To date, genome-wide association studies have identified at least 39 genes that are associated with statin treatment efficacy (Gryn and Hegele, 2014). Most of these genes are involved in either the direct pharmacokinetic handling of statins or in lipid metabolism pathways especially these involving cholesterol, the main target of statin therapy (Mangravite et al., 2006). However, accumulating evidence indicates that statins can also lower levels of triglycerides, potentially through altering degradation of apolipoprotein B (ApoB) and related very low-density lipoprotein (VLDL) balance, although the precise mechanism remains unclear (Ginsberg et al., 1987; Arad et al., 1992; Ginsberg, 1998).

One gene of particular interest within this context is *APOA5*, which has been repeatedly associated with the high inter-individual variations of serum triglycerides in all reported populations (Baum et al., 2003; Lai et al., 2004; Hubacek et al., 2008; Ouatou et al., 2014; Son et al., 2015) since its identification in 2001 (Pennacchio et al., 2001; van der Vliet et al., 2001). According to one estimation, the *APOA5* promoter SNP rs662799 (-1131T>C) alone can contribute to 6.2% of the genetic component of hypertriglyceridemia (Hegele, 2009). Of note, the minor C allele is much more common in the Asian population (26%-40%) than in Caucasians (only ~8%) (Baum et al., 2003). In addition, accumulating evidence suggests that this gene also confers risk for cardiovascular disease (Lai et al., 2004) and myocardial infarction (Do et al., 2015). Although previous studies have suggested a link between this gene and statin treatment (Brautbar et al., 2011; O’Brien et al., 2015), available statins differ in terms of their pharmacodynamic and pharmacogenetic properties (Kivisto et al., 2004; Schachter, 2005) and potency (Palmer et al., 2013; Arshad, 2014; Karlson et al., 2016). To the best of our knowledge, no well-designed prospective study, has investigated whether APOA5-statin interactions dependent on the statin type while controlling for differences in potency of the statins. One retrospective study did not observe an effect of statin type when investigating the interaction between the *APOA5* rs662799 variants and statins (Hubacek et al., 2009); however, this study did not include rosuvastatin, which is often considered to be a better treatment choice (Scott et al., 2004; McKenney, 2005; Schachter, 2005).

Here, we performed a pilot study to explore APOA5-statin interactions in 195 Chinese subjects randomized to rosuvastatin, atorvastatin, or simvastatin therapy for 12 weeks. To address whether the clinical responses of three types of statins differ between subjects with the same *APOA5* genetic background, we genotyped *APOA5* rs662799 SNP and measured the fasting plasma concentrations of triglycerides, cholesterols, free fatty acids, and four apolipoproteins both before and after statin treatments.

## Materials and Methods

### Study subjects and study design

We recruited 195 patients at Shanghai Ruijin Hospital Luwan Branch (affiliated to Shanghai Jiaotong University). In brief, the inclusion criteria were: (i) aged 18 years or older; (ii) newly diagnosed with dyslipidemia and/or increased risk of atherosclerotic cardiovascular disease and recommended to receive statins according to the 2013 American College of Cardiology (ACC) and the American Heart Association (AHA) Blood Cholesterol Guidelines (Stone et al., 2014); (iii) absence of major systematic diseases such as malignancy; and (iv) without medication (especially antibiotics) in the previous three months except antihypertensive therapy.

The subjects were then randomly divided into three treatment arms to receive rosuvastatin (5-10 mg per day), atorvastatin (10-20 mg per day), or simvastatin (40 mg per day) for 12 weeks. To achieve comparable clinical efficacies in response to the three statins, the different statin doses were selected based on both clinical practice and evidence suggesting that each rosuvastatin dose is equivalent to 3-3.5 times of atorvastatin and 7-8 times of simvastatin (at least in terms of cholesterol reduction) (Hubacek et al., 2009). All treatments were tolerated with no side effects reported.

Written informed consent was obtained from all the study participants. This study conforms to the ethical guidelines of the 1975 Declaration of Helsinki and was approved by Ethics Committee of Shanghai Ruijin Hospital Luwan Branch. Complete clinical trial registration is deposited at chictr.org.cn (ChiCTR-RRC-16010131).

### Laboratory analyses

Fasting plasma concentrations of triglycerides, total cholesterol, HDL cholesterol (HDLc), LDL cholesterol (LDLc), free fatty acids (FFA), and three different apolipoproteins (ApoA1, ApoB-100, ApoE) and lipoprotein(a) were measured by enzymatic methods using a Beckman Coulter Chemistry Analyzer AU5800 Series (United States) at both baseline and 12 weeks after treatments. ApoA5 was not measured since consistent evidence suggests rs662799 polymorphisms were not associated with the circulating levels of this apolipoprotein (Talmud et al., 2006; Henneman et al., 2007)

DNA was isolated using the TIANamp Blood DNA kit (purchased from Tiangen, Beijing, China) and individual *APOA5* variants (-1131T>C – rs662799) were genotyped using a base-quenched probe method combined with polymerase chain reaction (PCR) as described before (Luo et al., 2009). In brief, a 19-nt probe (5’-GGCAAATCTCACTTTCGCT-3’) containing the targeted SNP site was first conjugated with 6-carboxyfluorescein and then hybridized to its complementary target sequence from PCR amplification. An analytical melting program that involves heating the amplicon/probe heteroduplex will produce different fluorescence curves depending on the genotypes of rs662799. Both the probe and primers (forward: 5’-AGGAGTGTGGTAGAAAGACCTGTTG-3’; reverse: 5’-AACTACCCAGAGTCACTGTGTCCC-3’) used were synthesized by Sangon (Shanghai, China).

### Statistical analysis

Statistical differences between groups were estimated by Wilcox rank-sum test (between two groups), Kruskal-Wallis test (among three groups) for continuous variables or by Chi-square test for categorical variables. Different linear regression models were also built and compared using Chi-square test to confirm the effect of *APOA5* genotype on different biomarkers and adjusted for contributions from type 2 diabetes and sex. Associations between apolipoproteins and concentrations of LDLc and HDLc were measured by Spearman’s rank correlation analysis. Hardy-Weinberg Equilibrium was accessed by exact test based on R package “HardyWeinberg” (Graffelman, 2015). Raw P values were adjusted by the Benjamini-Hochberg method (Benjamini and Hochberg, 1995) with a false discovery rate of 5%. A power of 99.98% was obtained using pwr package (Champely, 2015) for this study based on 65 patients with paired design, 5% significance, and an estimated effect size of 0.7 for statin in reducing LDL cholesterol (Cholesterol Treatment Trialists et al., 2012; Ridker et al., 2016). All statistical tests and data visualizations as well as the stratified randomization process by considering BMI as covariate were performed under the R environment (Team, 2015).

## Results and discussion

### Baseline characteristics

The minor C allele frequency of *APOA5* rs662799 SNP in our cohort was 30%, consistent with other reports based on larger Chinese cohorts (Baum et al., 2003; Jiang et al., 2010); the genotype frequency of *APOA5* was in agreement with Hardy-Weinberg equilibrium (*n* = 13, 91 and 91 for C/C, T/C, and T/T allele carriers, respectively; *P*=0.171). T/C and C/C subjects were pooled as T(C)/C (*n* = 104) for further analyses to increase the power. With the exception of ApoE, there were no significant baseline differences between the treatment arms, including the frequencies of the T(C)/C and T/T genotypes (*P*=0.342) (**Table 1**). These data suggest that the treatment groups are in general homogeneous and this study design is suitable for addressing the relationship between *APOA5* variations and the clinical responses of three statins. When dividing the subjects by genotype, the T(C)/C allele carriers had significantly higher plasma triglycerides than T/T carriers at baseline (**Table 1**), in agreement with previous studies (Baum et al., 2003; Lai et al., 2004; Jiang et al., 2010).

**Table 1.**
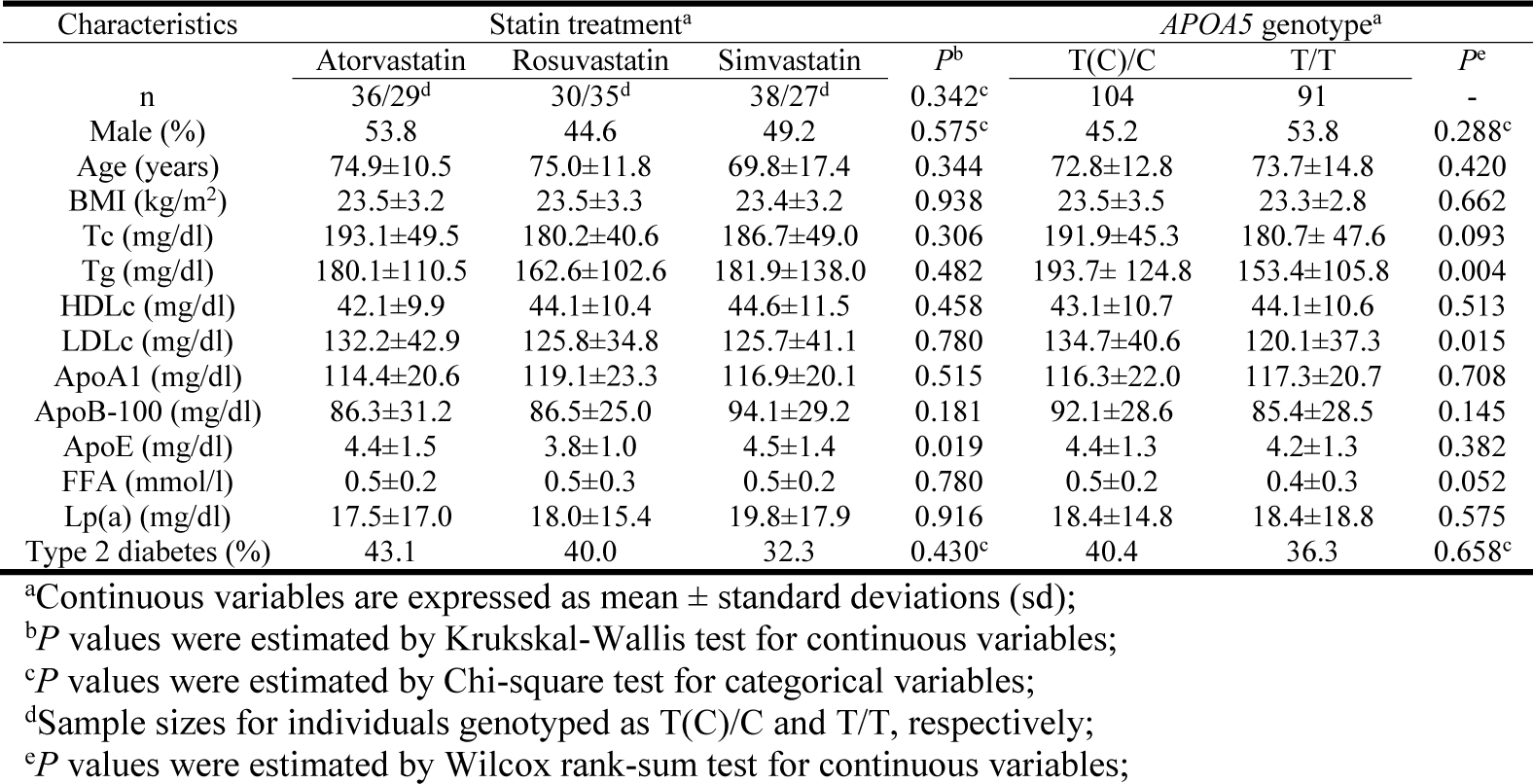
Baseline characteristics summarized by statin treatment and *AP0A5* genotypes, respectively.

We also noted that subjects with the T(C)/C genotype had higher LDLc than T/T carriers at baseline (**Table 1**); these findings were consistent with observations in a larger cohort (Lai et al., 2004) but an earlier study in Chinese men did not observe significant APOA5-LDLc interactions (Baum et al., 2003). It is not clear how *APOA5* variants affect LDLc as ApoA5 has only been detected on HDL and VLDL and not on LDL particles (Ballantyne et al., 2006). However, ApoA5 has been shown to directly interact with members of the LDL-receptor family (Nilsson et al., 2007). In addition, an earlier study has shown a significant association between the *APOA5* rs662799 SNP and increased risk of early-onset myocardial infarction even after adjusting for triglycerides (De Caterina et al., 2011), providing further evidence that this SNP may simultaneously affect other atherogenic lipids such as LDLc. It is also possible that this SNP is in complete linkage disequilibrium with other polymorphism(s) that can explain the observed LDLc levels.

### Rosuvastatin-induced changes in HDLc and triglycerides are not affected by APOA5 genotype

We next compared the clinical efficacies (in terms of cholesterol, triglyceride, and apolipoprotein changes) of the statins. As expected, all three statins promoted significant reductions in total cholesterol, ApoB, LDLc, ApoE and triglycerides and significant increases in ApoA1 and HDLc (**Figure 1A**). However, there were no significant differences between the treatment arms for any of the statin-induced changes in clinical biomarkers after adjusting for multiple testing (false discovery rate 5%), confirming that the response to 5-10 mg of rosuvastatin is similar to that of 10-20 mg atorvastatin and 40 mg of simvastatin as suggested previously (Hubacek et al., 2009). In agreement, results from a meta-analysis (Karlson et al., 2016), comparative pharmacology (McTaggart, 2003) and the MERCURY II clinical trial (Ballantyne et al., 2006) have all shown that rosuvastatin is more potent than the other statins and thus lower doses can be used to achieve equivalent responses.

**Figure 1.**
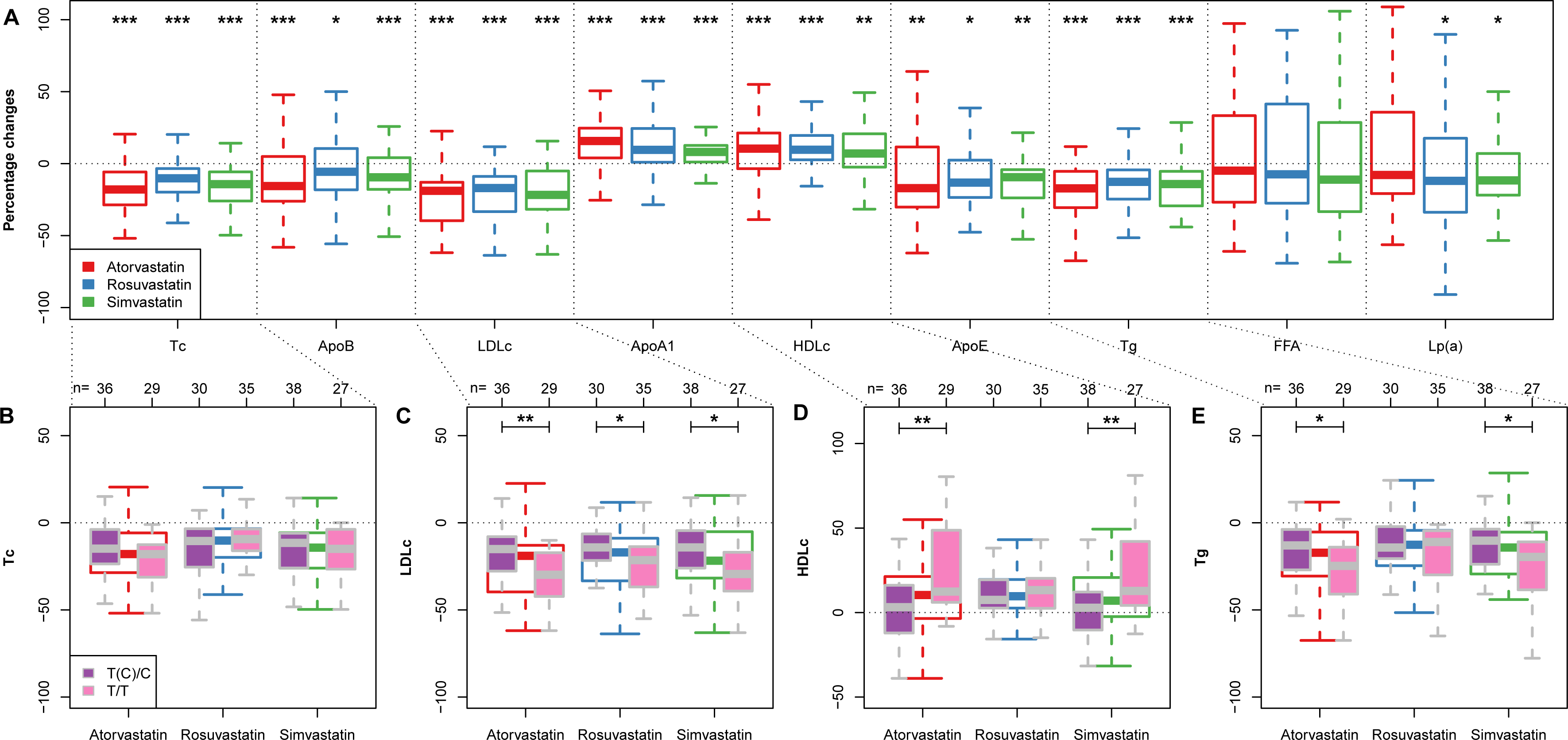
*APOA5*-statin interactions. (A) Box plots (with median) showing percentage changes in the indicated biomarkers after treatment with rosuvastatin (5-10 mg per day), atorvastatin (10-20 mg per day) or simvastatin (40 mg per day). *, *P*<0.05; **, *P*<0.01; ***, *P*<0.001 versus before treatment. (Wilcoxon signed-rank test) (B-E) Box plots (with median) showing percentage changes in total cholesterol (Tc) (B), LDLc (C), HDLc (D), and triglycerides (Tg) (E) in response to each statin in subjects divided by genotype (*APOA5* rs662799 T(C)/C and T/T). Sample sizes for each subgroup are given on top of panels B-E. *, *P*<0.05; **, *P*<0.01 (Wilcoxon rank-sum test).

To determine how *APOA5* variations affected the clinical responses of the three statins, we investigated how changes in the biomarker concentrations in response to each statin varied between subjects with the T(C)/C or T/T genotype (**Figure 1B-E; Supplementary Table S1).** Genotype did not affect the changes in total cholesterol (**Figure 1B**), apolipoproteins, FFA or lipoprotein(a) (data not shown) in response to any of the three statins. However, patients homozygous for the major T allele (T/T genotype) not only exhibited lower baseline LDLc levels (**Table 1**) but also demonstrated significantly larger LDLc reductions compared with the C allele carriers, independent of the type of statin used (**Figure 1C**). By contrast, rosuvastatin-induced changes in HDLc and triglycerides showed little variation between patients with the T(C)/C and T/T genotypes whereas changes in HDLc and triglycerides were more pronounced in T/T compared with T(C)/T carriers upon atorvastatin or simvastatin treatment (**Figure 1D,E**). These data suggest that rosuvastatin-induced responses may be less affected by *APOA5* variations than the other two statins. The results were still valid after adjusting for type 2 diabetes and sex **(Supplementary Table S2).** It has been suggested that the hydrophilic rosuvastatin is largely excreted unchanged (Martin et al., 2003) whereas the other two lipophilic statins undergo substantial metabolism by the CYP450 pathways and thus are more affected by gene polymorphisms (Kivisto et al., 2004; Schachter, 2005), consistent with our findings. Additionally, rosuvastatin differs from the other statins by its stronger binding to 3-hydroxy-3-methylglutaryl coenzyme A (HMG-CoA) reductase, lower systemic bioavailability, longer elimination half-life (McTaggart, 2003) and greater hepatoselectivity (Schachter, 2005). These physio-biochemical differences may also potentially contribute to the different treatment responses according to genotype. However, to fully understand how *APOA5* affects statin treatments, in-depth characterizations of its functional role are needed.

### Statin-APOA5 interactions altered the correlations between apolipoproteins and LDLc/HDLc

Although most therapies to reduce cardiovascular disease risk currently focus on reduction of LDLc and triglycerides, atherogenic proteins such as ApoB have also been suggested to have great predictive value (Ballantyne et al., 2008). Accordingly, the American Diabetes Association and the American College of Cardiology Foundation recommend that therapy for patients with high cardiovascular disease risk should aim to lower ApoB concentrations to below 90 mg/dl in addition to reducing LDLc levels (Brunzell et al., 2008). To address whether the well-known strong associations between ApoB and LDLc both before and after stain treatments (Ballantyne et al., 2008) differ among patients with different *APOA5* genotypes, we additionally analyzed ApoB-LDLc correlations within each *APOA5* SNP subgroup. Before treatment, strong and significant positive correlations were observed between ApoB and LDLc for both T(C)/C (Spearman coefficient rho= 0.55; *P*<0.001) and T/T carriers (rho=0.78; *P*<0.001; **Figure 2A**). After treatment, a comparable strong correlation only existed for the C allele carriers (rho=0.50; P<0.001; **Figure 2B**). In contrast, the dramatic decrease in the ApoB-LDLc correlation among T/T carriers (from 0.78 to 0.44) indicates the statin-induced reduction of ApoB in absolute values was much smaller than reduction of LDLc. Thus, further treatment to reduce the levels of ApoB even after achieving recommended LDLc reductions could be beneficial in T/T carriers. Similar observations were found between ApoA1 and HDLc **(Supplementary Figure S1).**

**Figure 2.**
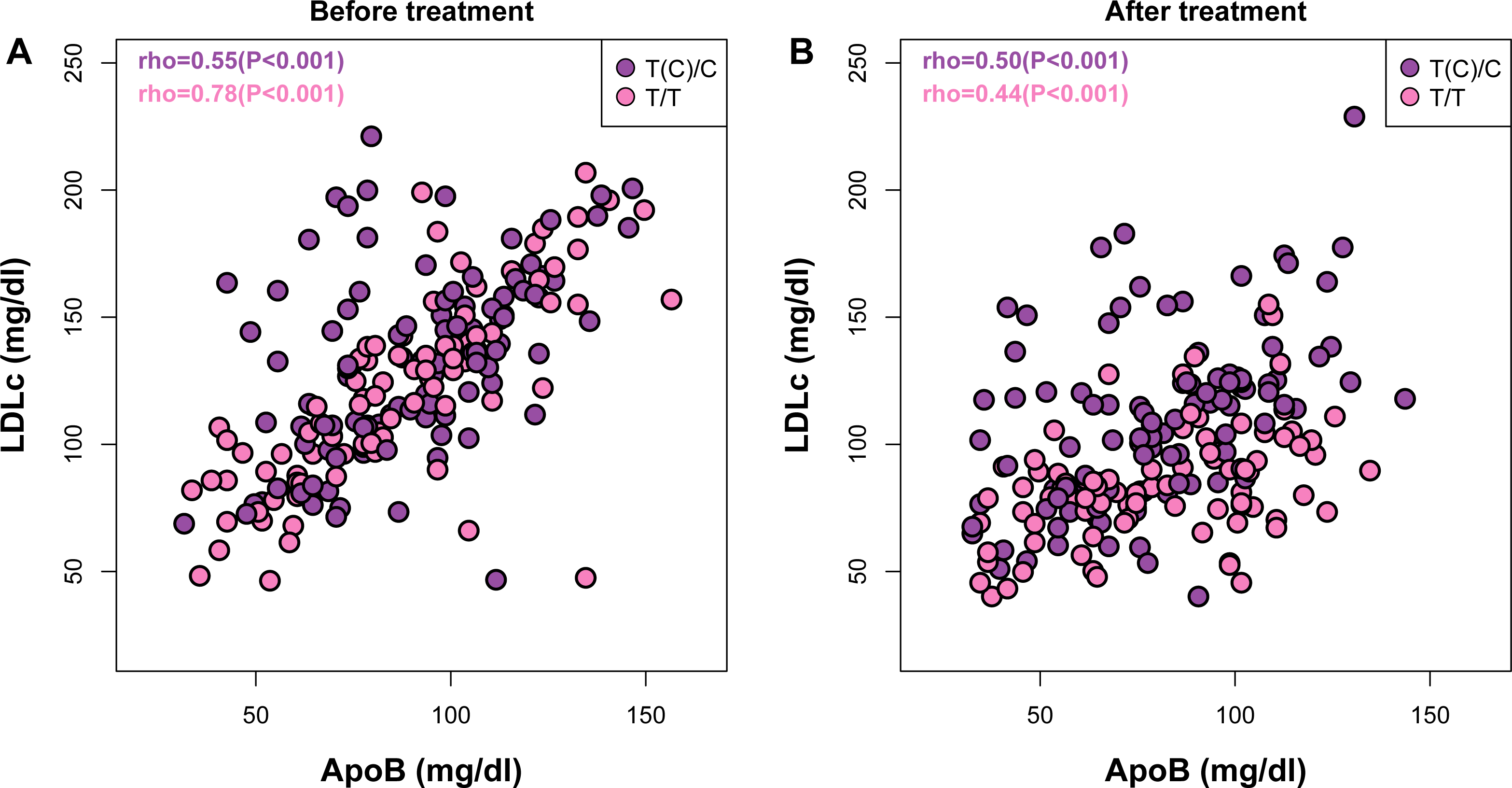
Both *APOA5* and statin alter the ApoB-LDLc correlations. Correlations between ApoB and LDLc before (A) and after (B) statin treatment in subjects with *APOA5* rs662799 T(C)/C or T/T allele.

## Conclusion

In summary, our results show that low-dose rosuvastatin achieves improvements in clinical responses that are comparable to those observed with higher doses of atorvastatin and simvastatin but are less affected by *APOA5* genotype. These findings support the growing recognition that rosuvastatin is a potentially better treatment option for patients with dyslipidemia and/or at high risk of cardiovascular diseases. In addition, integrated efforts, such as the NIH Pharmacogenetics Research Network (Giacomini et al., 2007), should be encouraged in the era of precision medicine to accelerate pharmacogenetics or pharmacogenomics research. Future studies should also consider stratifying populations by genetic background and by statin type.

## Author contributions

S.H., G.L., N.X. and J.Z. designed the study; S.H. performed the randomization process and clinical intervention; J.L. and W.W. enrolled participants and measured the lipids and apolipoproteins; J.Z. and G.L. performed the genotyping analysis; S.H., C.M., J.L. and W.W. collected and analyzed the data; S.H., C.M, G.L. and J.Z. wrote the manuscript.

## Acknowledgements

The authors would like to thank Dr. Hao Wu and Prof. Jan Borén from University of Gothenburg as well as Dr. Bingxiang Xu from Beijing Institute of Genomics for help discussions on the manuscript and data analysis. We also appreciate the great help from Dr. Rosie Perkins for editing our manuscript. This study was supported by a funding to S.H. from the Science Committee Foundation of Huangpu District, Shanghai, China (HKW201503).

